# Toward a single dose cure for Buruli ulcer

**DOI:** 10.1101/2020.04.15.044115

**Authors:** Sangeeta S Thomas, Nitin Pal Kalia, Marie-Thérèse Ruf, Gerd Pluschke, Kevin Pethe

**Author notes:** **Corresponding author:** Kevin Pethe, Lee Kong Chian School of Medicine, Nanyang, Technological University, 11 Mandalay Road, Singapore, 308232.

## Abstract

A single dose of TELACEBEC (Q203), a phase 2 clinical candidate for tuberculosis, eradicates *Mycobacterium ulcerans* in a mouse model of Buruli ulcer infection without relapse up to 19 weeks post treatment. Clinical use of Q203 could dramatically simplify the clinical management of Buruli ulcer, a neglected mycobacterial disease.

## Text

Buruli ulcer is a chronic ulcerating disease of the skin and underlying tissues caused by *Mycobacterium ulcerans*. The disease is regaining importance in West Africa and South East Australia with increasing incidence and severity (1, 2). The current treatment strategy involves an eight-week regimen of rifampicin administered with streptomycin or clarithromycin (3, 4). Disease management is complicated by an underreporting, especially in rural Africa (5), social stigmata, and lack of awareness that impede the deployment of medical treatment. Inadequate therapy may drive a substantial number of permanent disabilities, especially in children. Compliance to an eight-week therapy is also a serious limitation.

Telacebec (Q203) is an imidazopyridine amide drug targeting the mycobacterial cytochrome *bcc:aa_3_* terminal oxidase. The drug candidate, currently in clinical trial phase 2 for tuberculosis (6), has excellent activity against *M. ulcerans in vitro* and *in vivo (7–9)*. In *Mycobacterium tuberculosis*, the bactericidal potency of Q203 is limited by the presence of the cytochrome *bd* oxidase, an alternate terminal oxidase. The exquisite sensitivity of *M. ulcerans* to Q203 is explained by the absence of a functional cytochrome *bd* oxidase in this species (10, 11).

Considering the distinct potency of Q203 coupled with a long half-life and favourable toxicological profile (7, 10), we evaluated the potency of a single dose of Q203 to eradicate *M. ulcerans* in an established mouse model of Buruli ulcer infection. BALB/c mice were infected in the left hind footpad with 1.1 × 10^5^ colony-forming units of *M. ulcerans* S1013 as described by Fenner *et al* (12). Disease progression was monitored by weekly measurements of footpad thickness (Figure 1A). Treatment was initiated 5 weeks post-infection when the mean footpad swelling reached 3.7 mm, reflecting the establishment and progression of the infection. Fourteen animals were randomly assigned to the following treatment categories: rifampicin (10 mg/kg) + clarithromycin (100 mg/kg) administered 5 times per week for 4 weeks (20 total doses); Q203 at 20 mg/kg administered only once (single dose), Q203 at 5 mg/kg administered weekly for 4 weeks (4 doses); and bedaquiline at 20 mg/kg administered only once (single dose). The diarylquinoline bedaquiline (Sirturo®) was selected as a comparator because the drug acts on the Oxidative Phosphorylation pathway as well (13), shares with Q203 a very long half-life (13), and has a demonstrated potency against *M. ulcerans* (14) . All drugs were administered orally.

**Figure 1.**
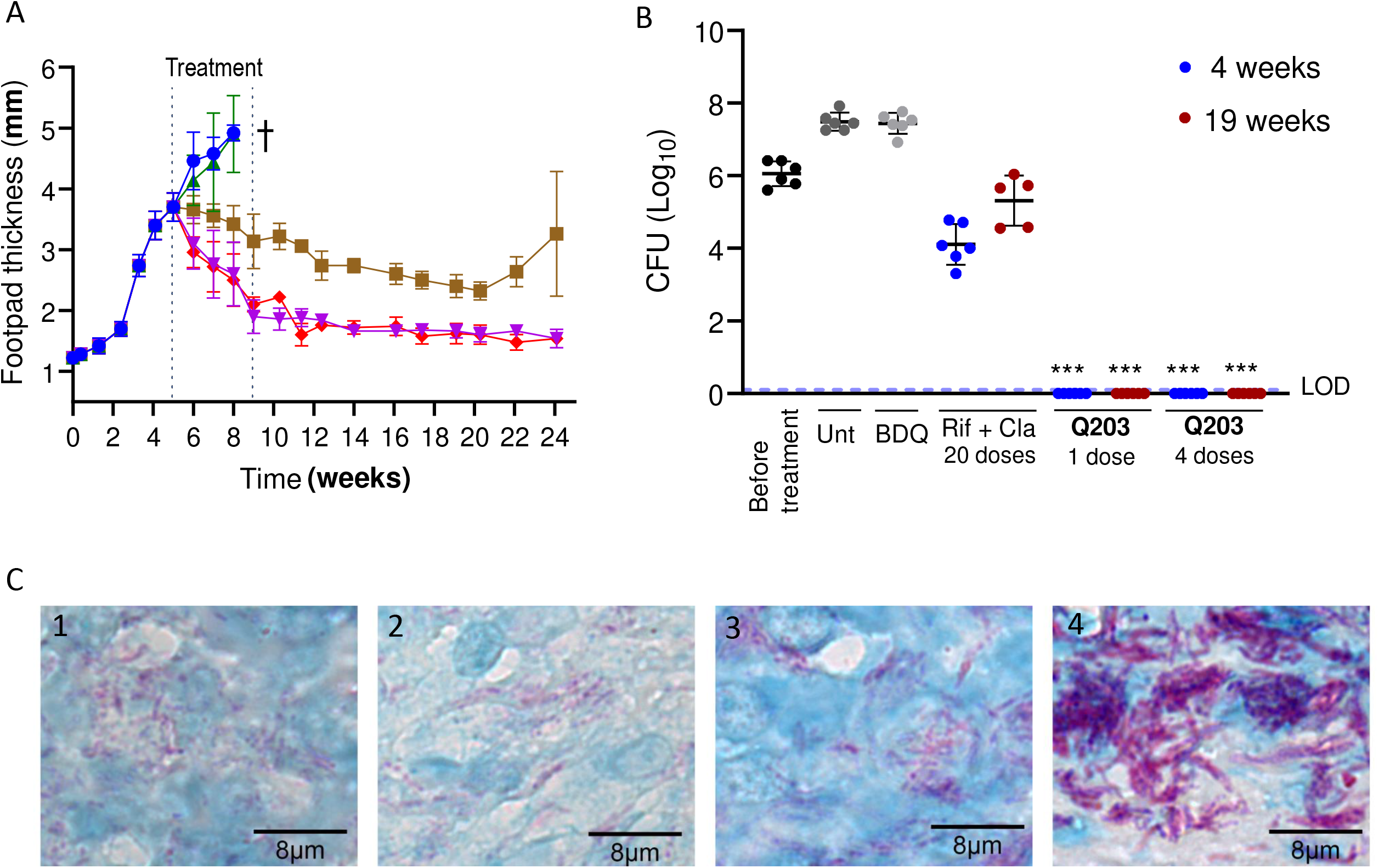
A single dose of Telacebec (Q203) is curative in a mouse model of Buruli ulcer. A) Mice were infected with 1.1 × 10^5^ CFUs of *M. ulcerans* S1013 five weeks prior treatment initiation. Mice were randomly assigned to oral treatment with either 1 dose of Q203 at 20 mg/kg (red diamonds), 4 doses of Q203 at 5 mg/kg (purple inversed triangles), 1 dose of bedaquiline at 20 mg/kg (BDQ, green triangles), 20 doses of Rifampicin at 10 mg/kg + clarithromycin at 100 mg/kg (brown squares), or dosing vehicle alone (Unt., blue circles). Footpad thickness was measured with a calliper weekly over 24 weeks period. Untreated mice (n=9) and mice treated with BDQ (n=14) were euthanised at week 8 post infection due to unfavourable disease progression (indicated by a cross); B) Bacterial loads in the infected feet were enumerated by CFU count on agar plates at the indicated time points, except for the untreated group (week 5 and week 8) and the BDQ-treated group (week 8). Six animals per time point were used; statistical analysis was performed using Student t-test. *** P value<0.0001, when comparing CFU of Q203 (1 and 4 doses) to Rifampicin + Clarithromycin at corresponding 4 and 19 weeks; C) Histopathological analysis of infected footpads after Ziehl–Neelsen / Methylene blue staining. Note the faintly stained acid-fast debris and the absence of intact Acid-fast Bacillus (AFBs) in the animals treated with 1 dose (panel 1) and 4 doses (panel 2) of Q203. The structure of some of the bacilli was more preserved in animals treated for four weeks with Rifampicin + Clarithromycin (panel 3), while numerous clusters of solid-stained AFBs were present in the untreated footpads at week 8 (panel 4). The animal protocol used in this study was approved by the Institutional Animal Care and Use Committee of Nanyang Technological University (Protocol # A18022).

In the untreated group, the disease progressed steadily as witnessed by an increase in feet inflammation and swelling. At 8 weeks post-infection, the feet thickness reached 4.9 mm (Fig. 1A) and some of the lesions stared to ulcerate. For these reasons, the animals had to be euthanized. In comparison, the disease progression stopped rapidly in the animal treated with Q203; the feet pathology reversed as early as one week post-treatment (Fig.1A). In comparison, complete reversal of swelling was not achieved in the animals treated with 20 doses of rifampicin + clarithromycin. The swelling of the feet remained consistently higher in the rifampicin + clarithromycin compared to the Q203 groups throughout the observation period (Fig.1A). The high potency of a single and weekly (4 doses) administration of Q203 was confirmed by Colony Forming Units (CFU) counts in the infected feet. The bacterial load diminished rapidly in the animal treated with Q203, reaching the limit of detection (1 CFU) at 4 weeks post-treatment. No relapse was observed up to 24 months post-infection (Fig.1B). Conversely, under similar conditions, *M. ulcerans* bacilli were detected in all the animals treated by the combination Rifampicin + clarithromycin (average CFU: 2.3 × 10^4^, range: 6 × 10^3^ - 6 x 10^4^, Fig.1B) at 4 weeks post-treatment.

Histopathology analysis supported the curative potency of a single dose of Q203. Only faintly stained acid-fast debris, but no intact acid-fast bacilli (AFB) were observed at week 9 in the animals treated with a single dose (Fig.1C1) or four doses of Q203 (Fig.1C2). In animals treated for four weeks with 20 doses of Rifampicin + Clarithromycin, the structure of some of the bacilli was more preserved, suggesting incomplete killing (Fig.1C3), confirming the CFU count on agar plates. In contrast, numerous clusters of solid-stained AFB were present in the untreated footpads at week 8 (Fig.1C4)(15).

The feet swelling of the animals treated for four weeks with the rifampicin + clarithromycin combination diminished steadily until week 15 post-treatment, but increased again thereafter, suggesting a relapse that was confirmed by an increase in the CFU at week 19 post-treatment (Average CFU: 4.3 × 10^5^, range: 3.6 × 10^4^ – 1 × 10^6^) (Fig.1A,B). This observation confirms that a 4-week regimen of this drug combination is insufficient to clear the infection (3). It was interesting to note that despite a favourable pharmacokinetic profile, a single dose of bedaquiline was ineffective at controlling disease progression (Fig.1A,B). The lack of potency at a single dose of bedaquiline is probably linked to its relatively modest potency compared to Q203 (10). Newer promising diarylquinolines with improved potency and favourable pharmacokinetic parameters should be explored for potency against *M. ulcerans*.

Q203 holds exceptional promises as a drug candidate for Buruli ulcer. The role of the drug candidate in abbreviating therapy to two weeks has been recognised in the mouse model (9). The proof of concept of a single dose cure opens the possibility to develop a drastically simplified regimen to treat the disease. Future combination studies between Q203 and optimized diarylquinolones (16) should be performed to develop a single dose combination cure with minimum risks for emergence of resistant mutants.

## Acknowledgements

This work was supported in part by the Singapore Ministry of Health’s National Medical Research Council under its Cooperative Basic Research Grant (Project Award NMRC/CBRG/0083/2015), the Lee Kong Chian School of Medicine, Nanyang Technological University Start-Up Grant (K.P.), and the National Research Foundation Competitive Research Programme (CRP), Grant Award Number NRF–CRP18–2017–01. S.S.T. was supported by an Interdisciplinary Graduate School Scholarship from the Nanyang Technological University. We thank Michael Berney for critical reading of the manuscript.

